# Lysine methyltransferase SETD6 modifies histones and non-histone proteins

**DOI:** 10.1101/092031

**Authors:** Olivier Binda

## Abstract

Although central to regulating the access to genetic information, most lysine methyltransferases remain poorly characterised relative to other family of enzymes. Herein, we report new substrates for the lysine methyltransferase SETD6. Based on the SETD6- catalysed site on the histone variant H2AZ, we identified similar sequences in the canonical histones H2A, H3, and H4 that are modified by SETD6 *in vitro*, and putative non-histone substrates. We herein expend the repertoire of substrates for methylation by SETD6.

## INTRODUCTION

Histone H3 lysine residues were found to be methylated over fifty years ago*^(1, 2)^*. However, it was not until 2000 that the first mammalian histone lysine methyltransferase was discovered*^(3)^*. The latest human genome annotation predicts over 60 lysine methyltransferases (KMTs) based on sequence similarities with either a SET or a seven β-strand catalytic domain*^(4)^*. However, most of these enzymes remain uncharacterized or poorly studied. Thus, important questions regarding the biological relevance and biochemical properties of these enzymes remain unanswered. Importantly, several histone KMTs also methylate non-histone substrates, such as the tumour suppressors p53*^(5–7)^*, ING2*^(8)^*, and pRB*^(9)^*, as well as chromatin proteins such as DNMT1*^(10)^*, HP1α/β/γ*^(11)^*, RUVBL1 ^*(12)*^, and RUVBL2 *^(13)^*.

The methyltransferase SETD6 mono-methylates the NFκB subunit RelA at lysine 310 (K310^me1^)*^(14)^*, the histone variant H2AZ at lysine 7 (K7^me1^)*^(15)^*, and the kinase PAK4*^(16)^*. The expression of SETD6 is amplified in about 10% of cases of breast cancer according to a study using a patient xenograft model*^(17)^* and is required for cellular proliferation in both ER^+^ and ER^−^ breast cancer cell models*^(18)^*, suggesting an important role in driving breast cancer progression. Indeed, SETD6 was recently found to associate with the cytoskeleton protein VIM*^(19)^*, which is involved in epithelial to mesenchymal transition (EMT), cellular attachment, migration, and signalling, suggesting a role in metastasis.

Much like classical signal transduction events involve phospho-dependent protein-protein binding, chromatin signalling events implicate post-translational modifications in the regulation of macro-molecules interactions. For example, lysine methylation of histones lysine residues serves as landing pads for chromatin proteins, which are referred to as histone mark readers or simply readers, thereby nucleating enzymatic complexes that modify and remodel chromatin to regulate access to genetic information.

Herein, we demonstrate that recombinant SETD6 methylates canonical histones H2A, H3, and H4, as well as linker histones H1 and the non-histone protein ING2 *in vitro*, and identify several putative novel substrates, including chromatin proteins and other lysine methyltransferases.

## RESULTS

We previously identified 2 mono-methylation sites on the histone variant H2AZ catalysed by SETD6*^(15)^*. Interestingly, these modified sites, H2AZK4^me1^ and H2AZK7^me1^, are similar. Both modified lysine residues are preceded by a small amino acid (alanine or glycine) at position −2 and a glycine at position −1 (**Table 1**). Examination of canonical histone tails revealed similar sequences in histones H2A, H3, and H4, and identification of a putative SETD6 methylation consensus motif A/G/RGK^me1^A/GG (**Table 1**).

**Table 1:** SETD6 consensus motif. Based on H2AZ methylation sites by SETD6, putative modification sites were identified in canonical histones H2A, H3, and H4.

| Histone | Consensus |  |  |  |  |  |  |
| --- | --- | --- | --- | --- | --- | --- | --- |
|  | -3 | -2 | -1 | 0 | +1 | +2 | +3 |
| H2AZ | A | G | G | K<sup>4</sup> | A | G | K |
| H2AZ | K | A | G | K<sup>7</sup> | D | S | G |
| H2A | G | R | G | K<sup>5</sup> | Q | G | G |
| H3 | T | G | G | K<sup>14</sup> | A | P | R |
| H4 | G | R | G | K<sup>5</sup> | G | G | K |
| H4 | G | L | G | K<sup>12</sup> | G | G | A |
|  | . | . | * | * | . | : | . |

To test whether SETD6 could methylate these other histones, we used a mixture of purified calf thymus histones as substrates. Interestingly, SETD6 was capable of modifying the linker H1 histones as well as the canonical histones H2A, H3, and H4 (**Figure 1**). As a positive control, we used SET7, which is known to methylate H3*^(20)^*, and H1 histones*^(21)^*. As a negative control, GST alone was used and as expected GST had no detectable methyltransferase activity on histones (**Figure 1**).

**Figure 1:**
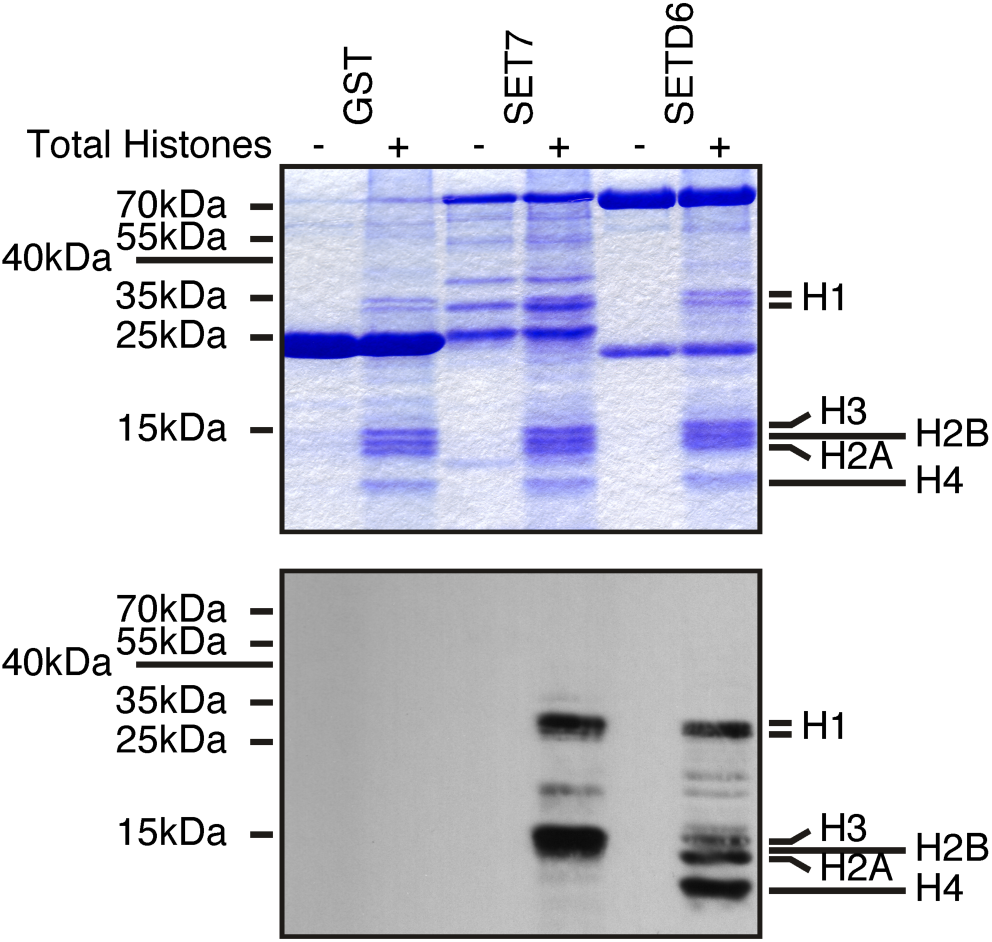
SETD6 methylates histones. Recombinant KMTs were used to modify histones isolated from calf thymus in the presence of ^3^H-SAM. Samples were analysed by SDS-PAGE, then either stained with Coomassie (top panel) or transferred to PVDF membrane and autoradiographed (bottom panel).

To confirm the SETD6-catalysed methylation sites on canonical histones, the first 50 amino acid residues of H2A, H3, and H4 were fused to the amino terminus of GST to leave the histone tail free at the amino terminus and generate H2A_1-50_-GST, H3_1-50_-GST, and H4_1-50_-GST. Then the predicted sites (**Table 1**) were converted to arginine by site-directed mutagenesis. The affinity purified recombinant proteins were then used for *in vitro* KMT assays with SETD6. In agreement with previous experiments showing that SETD6 modifies canonical histones (**Figure 1**), SETD6 methylated recombinant histone tails from H2A, H3, and H4 (**Figure 2A**). Importantly, mutation of the GK motifs reduced drastically the methylation of H2A and H3 by SETD6 (**Figure 2A**). However, single mutation of H4 at K5 or K12 only minimally reduced methylation by SETD6 (**Figure 2A**).

**Figure 2:**
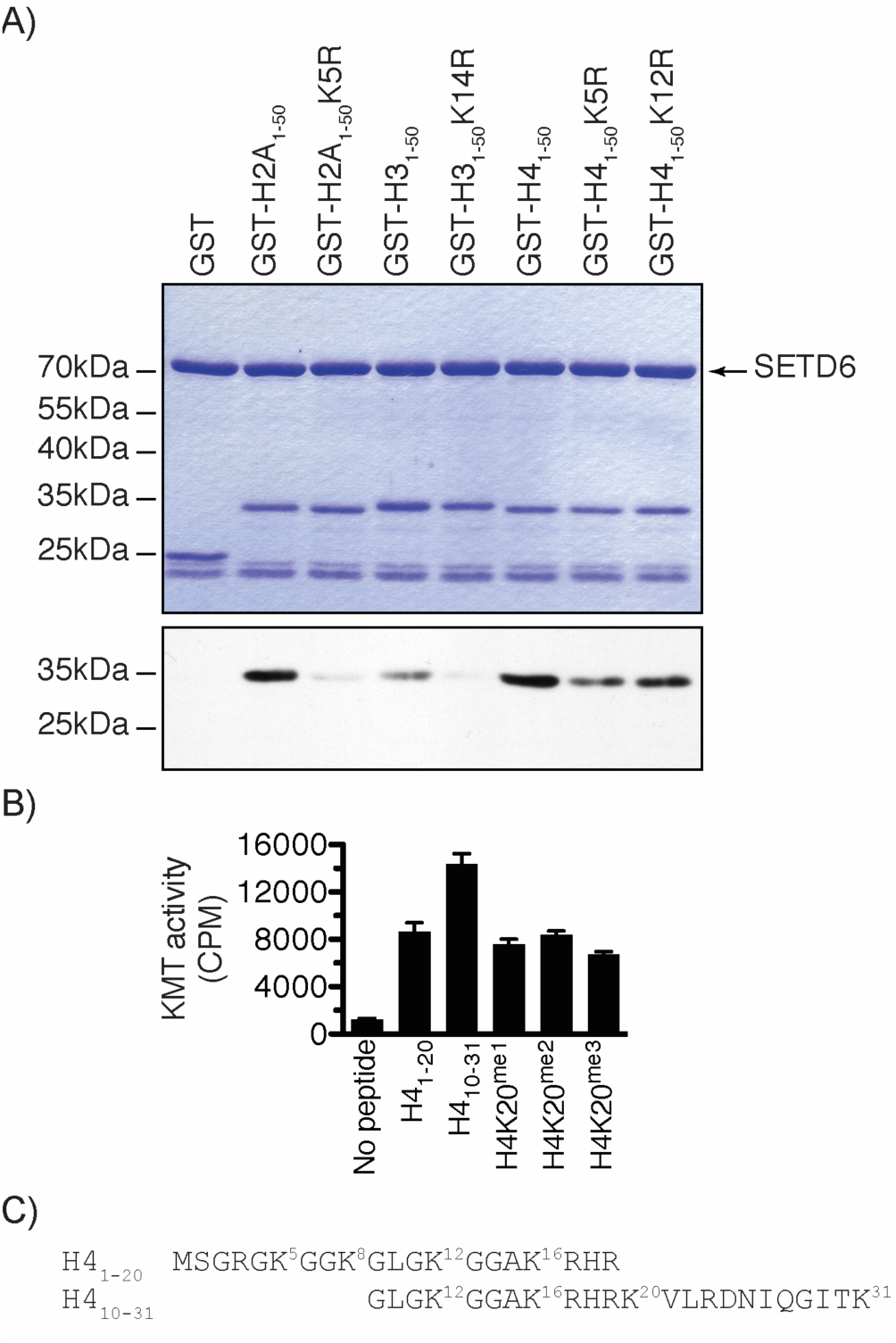
SETD6 methylates GK motifs in canonical histones. (A) SETD6 was used to methylate the indicated recombinant histone tails. Samples were analysed by SDS-PAGE, then either stained with Coomassie (top panel) or transferred to PVDF membrane and autoradiographed (bottom panel). (B) KMT assays were performed with SETD6 on H4 peptides. (C) Sequence of H4 peptides used in panel B with each lysine numbered.

Since H4 methylation seemed stronger (**Figure 1**), and to further investigate the methylation of H4 by SETD6, methyltransferase assays on H4 peptides H4_1-20_ and H4_10-31_ were performed and confirmed that SETD6 methylates H4 (**Figure 2B**). Interestingly, SETD6 methylated H4_10-31_ better than the H4_1-20_ peptide, while methylation at K20 (K20^me1^, K20^me2^, or K20^me3^) impaired this effect (**Figure 2B**), suggesting that SETD6 modifies H4K20 in addition to H4K12 or other site(s), such as H4K16 or H4K31 (**Figure 2C**).

Alternatively, these results may suggest that there is a cross-talk between H4K20^me^ and the SETD6-catalysed methylation site(s).

Several KMT, including SETD6, modify non-histone proteins. We thus searched for the GKDS motif in protein sequence repositories and identified several putative SETD6 substrates (**Table 2** and **S1**), including the ATPase RUVBL1, which is modified by the H3K9 methyltransferases G9A and GLP*^(12)^*. Importantly, some putative SETD6 substrates (AHNAK2, ERICH3, and MDN1) were found in the PhosphoSitePlus mass spectrometric database to be methylated at the predicted site*^(22)^* (**Table S1**). A similar search using the H4K5 and H4K12 motif GKGG also yielded several putative substrates for SETD6, such as the chromatin remodeller BRG1 (GK^1029^GG) and the HBO1 acetyltransferase subunit JADE2 (GK^638^GG).

**Table 2:** Novel SETD6 putative substrates. The GKDS motif from H2AZ was used to identify novel substrates for SETD6.

| Substrate | Site | Function |
| --- | --- | --- |
| AHR | GK<sup>438</sup> DS | Nuclear receptor |
| APAF1 | GK<sup>100</sup> DS | Apoptosis |
| CDKN2C | GK<sup>160</sup> DS | CDK4 inhibitor p18<sup>INK4C</sup> |
| GREB1 | GK<sup>278</sup> DS | Proliferation |
| KDM5A | GK<sup>1504</sup> DS | Demethylase |
| MYST4 | GK<sup>393</sup> DS | Acetyltransferase |
| RBBP8 | GK<sup>426</sup> DS | pRB-binding protein |
| RUVBL1 | GK<sup>422</sup> DS | Chromatin remodelling |
| SMYD4 | GK<sup>63</sup> DS | Putative KMT |
| SUV420H1 | GK<sup>864</sup> DS | H4K20 KMT |

Herein, we found that SETD6 methylates histones H2A, H3, and H4 on lysine residues within an A/G/ RGK^me1^A/GG consensus motif.

## DISCUSSION

Proteomic studies have identified post-translational modifications on histones and non-histone proteins, but the enzymatic activities depositing these modifications remain largely unknown. There is a dire need to identify PTMs to understand how proteins are regulated, but more importantly to identify the enzymes catalysing these biochemical events. To this end, we have in the past designed an unbiased chemical-biology approach to tag novel KMT substrates*^(23)^*. However, traditional biochemical studies are still required to investigate and validate novel post-translational modifications.

The H3K14^me1^ mark was reported to occur in both human and mouse*^(24, 25)^*, supporting the existence of the modification in cells. We have herein identified the first KMT capable of modifying this site *in vitro*. Further work will be required to validate the role of SETD6 in the catalysis of H3K14^me1^ in cells and the function of this mark.

Interestingly, H4K12^me3 *(22)*^, H4K16^me3^*^(22, 24)^*, and H4K31^me1 *(26)*^ were detected by mass spectrometry. However, no other reference to these modifications appear in the current literature. H4K5 was recently found to be methylated by SMYD3*^(27)^*.

Although mono-methylation events on the linker histone H1 H1F0 was reported at K12, K59, K82, K102, K108, and K155*^(22, 28)^*, these sites do not share similarities with the SETD6 consensus motif, suggesting that SETD6 modifies non-GKDS sequences. Indeed, SETD6 methylates RelA at the non-GKDS site FK310SI*^(14)^*. In addition, SETD6 can methylate the histone mark reader ING2 *in vitro* (**Figure S1**) and PAK4*^(16)^*, both of which do not contain any GKDS-like motif.

Together, the *in vitro* data provided here identifies SETD6 as a likely candidate for the methylation of reported events on H3K14, H4K5, H4K12, H4K16, and H4K31. In addition, we identified several putative non-histone protein substrates for SETD6.

## METHODS

### Plasmids

The modified pGEX plasmid with an engineered multi-cloning site at the N-terminus of the GST coding sequence was described previously*^(15)^*. The cDNA of histones H2A, H3, and H4 was amplified by PCR from reverse transcribed total RNA and inserted in frame with GST using restriction endonucleases and T4 DNA ligase (NEB).

### Recombinant protein expression and purification

Essentially, BL21 DE3 competent bacteria (Stratagene) were transformed with pGEX plasmids. Single colonies were picked and grown in 2YT media. Expression of GST-fusion proteins was induced with 0. 01mM IPTG for 2.5 hours at 37°C, cells were collected and lysed in buffer (50mM Tris-Cl pH 7.5, 150mM NaCl, 0.05% NP-40). Recombinant protein were batch purified using glutathione-sepharose beads. GST-SETD6 was purified similarly, but from Sf9 insect cells as described*^(15)^*.

### *In vitro* KMT and Flashplate KMT assays

Lysine methylation assays were performed in reaction buffer (50mM Tris-Cl pH8.0, 10% glycerol, 20mM KCl, 5mM MgCl2) supplemented with ^3^H-SAM as described*^(8)^*, using calf thymus histones (Worthington), recombinant histone tails (see above), or biotinylated histone H4 peptides.

### Motif search

The GKDS sequence was used in a motif search using PHI-BLAST against the Non-redundant protein sequences (nr) database, restricted to *Homo sapiens* (taxid:9606).

## ACKNOWLEDGMENTS

OB is supported by the Newcastle’s Biomedical Fellowship Programme, which is in part funded through the Wellcome Trust’s Institutional Strategic Support Fund, and by the Breast Cancer Campaign charity grant number 2013MaySP005.

